# El Niño, capture success, and wing fluctuating asymmetry in *Drosophila sturtevanti* Duda, 1927

**DOI:** 10.64898/2026.10.09.757947

**Authors:** Pedro Gabriel Carneiro Moreira, Caio de Castro Mello, Layla Patrícia Teixeira, Karla Yotoko

## Abstract

Fluctuating asymmetry (FA) is widely used as a proxy for developmental instability, although reduced rather than increased FA, has occasionally been reported under particularly challenging conditions, possibly because environmental filtering removes less developmentally stable individuals. Here, we investigated temporal variation in wing shape asymmetry in males of *Drosophila sturtevanti* collected in 2022, 2024, and 2026 at the same sampling site, an urban Atlantic Forest fragment. Sampling in 2022 and 2026 were completed during single field expeditions in early April, whereas in 2024 repeated collections were required until mid-May because of markedly low capture success. We therefore examined whether this difference was associated with unusual environmental conditions and whether FA was higher in 2024, as expected under greater developmental perturbation. Daily temperature and relative humidity were examined from early March onward, encompassing approximately one month before adult emergence. The 2024 sampling period was warmer and drier than the corresponding periods in 2022 and 2026 and coincided with the final phase of the strong 2023–2024 El Niño event. Contrary to our prediction, FA was significantly lower in 2024. The magnitude of directional asymmetry remained similar among years, although the spatial pattern of the DA vector differed between 2024 and 2026. The coincidence of reduced capture success and reduced FA is consistent with the hypothesis that adverse conditions may impose demographic filtering, such that individuals expressing greater developmental instability are less likely to contribute to the sampled adult population.

## 1. Introduction

More than a century ago, D’Arcy Thompson emphasized the central role of symmetry in the study of biological form and growth (Thompson, 1994; originally published in 1917). He used simple symmetrical forms to illustrate how organismal shape reflects the forces acting during growth, arguing that departures from symmetry necessarily imply some departure from uniformity in the underlying system. In bilaterally symmetrical organisms, corresponding structures on the left and right sides provide a particularly informative system for studying departures from symmetry because they share the same genotype and develop under largely equivalent external conditions (Klingenberg and McIntyre, 1998). Small differences between sides have therefore long been used to investigate variation in developmental processes, initially through measurements of individual morphological traits and, more recently, through geometric morphometric approaches that quantify asymmetry in the shape of entire structures (Klingenberg, 2022; Klingenberg & Mcintyre, 1998; Palmer & Strobeck, 1986).

Bilateral asymmetry can be decomposed into directional asymmetry (DA), in which one side differs consistently from the other at the population level, and fluctuating asymmetry (FA), which represents individual departures from the mean right–left difference (Zelditch *et al*., 2012). DA has traditionally been regarded as a relatively stable characteristic of populations, species, or broader taxonomic groups, reflecting consistent patterns of lateralized development (Klingenberg and McIntyre, 1998). In contrast, although the relationship between FA and developmental instability remains debated (e.g. Graham, 2021), FA has been widely used as a proxy for developmental instability (Beasley *et al*., 2013), based on the expectation that developmental perturbations may increase random departures from bilateral symmetry. Such perturbations may arise from environmental stressors, including nutritional conditions, temperature, chemical pollution, and population density (Arce et al., 2023; Benítez et al., 2020, 2025; Beasley et al., 2013), reduced heterozygosity or inbreeding (Soulé, 1967; Palmer & Strobeck, 1986), experimental host shifts (Soto et al., 2008; Bouzas et al., 2025), or other conditions that interfere with development.

Since 2015, our laboratory has studied a population of *Drosophila sturtevanti* Duda, 1927 (Diptera: Drosophilidae) from an urban Atlantic Forest fragment in southeastern Brazil. Repeated field surveys indicated that early April is a particularly suitable period for sampling this population. In 2022, we initiated geometric morphometric studies and found that laboratory maintenance altered wing shape as early as the first generation (Dias *et al*., 2026), prompting a new field collection in 2024. However, no *D. sturtevanti* were collected on the first 2024 sampling occasion, requiring repeated sampling until mid-May. In contrast, the species was readily collected during a single sampling occasion in April 2026.

These observations raised two related questions. First, could unusual environmental conditions have contributed to the low sampling success observed in 2024? Second, were differences among sampling years accompanied by changes in wing asymmetry? We therefore characterized temperature and relative humidity during the periods preceding and encompassing field sampling and quantified wing asymmetry in males collected in 2022, 2024, and 2026. We decomposed total asymmetry into directional and fluctuating components, expecting DA to remain relatively stable through time and FA to increase under the more challenging developmental conditions hypothesized for 2024.

## 2. Methods

### 2.1. Sampling

Field sampling was conducted in 2022, 2024, and 2026 at the same Atlantic Forest fragment in Minas Gerais, southeastern Brazil (20°45′48″ S, 42°51′53″ W), corresponding to the UFV-1 sampling site described in Moreira et al. (2026). On each sampling occasion, six banana-baited traps (Medeiros and Klaczko, 1999) were deployed for 48 h. After collection, specimens were transported to the laboratory, anaesthetized with carbon dioxide, sorted, and stored frozen. Male specimens used for geometric morphometric analyses were identified by examination of the aedeagus following Magalhães (1962).

### 2.2. Field environmental conditions

Because the adults collected in the field had developed under environmental conditions preceding the sampling dates, we characterized the climatic conditions during the period encompassing approximately one month before the usual sampling period. Under laboratory conditions, the life cycle of *Drosophila sturtevanti* from egg to adult takes approximately three weeks (Galego and Carareto, 2005)although developmental time under natural conditions is unknown. We therefore considered the interval from 1 March to 15 May in each sampling year (2022, 2024, and 2026), which encompasses the month preceding the usual early-April sampling period as well as the extended sampling period required in 2024.

Daily environmental data for the sampling locality were obtained from the NASA POWER database (https://power.larc.nasa.gov/) via its daily API, using the geographic coordinates of the sampling site (20°45′36.2″ S, 42°52′14.6″ W) for the period from 1 March to 15 May in each sampling year. Daily mean air temperature and relative humidity were represented by the T2M and RH2M variables, respectively. Data were accessed in R using `jsonlite` (Ooms, 2014), processed with `dplyr` (Wickham *et al*., 2026), and visualized using `ggplot2` (Wickham, 2016) and patchwork (Pedersen, 2019). These data represent gridded environmental estimates rather than measurements from a local meteorological station, but provide a consistent basis for comparisons among sampling years.

### 2.3. Wing preparation and landmark digitization

Male *Drosophila sturtevanti* collected in the field in 2022, 2024, and 2026 were used for geometric morphometric analyses. Only individuals with both wings sufficiently preserved for the identification of all anatomical landmarks were retained.

The wings were removed from frozen specimens at their insertion into the thorax and positioned together on a microscope slide in anatomical orientation. Both wings of each individual were photographed in the same image, together with a 1-mm scale, ensuring that left and right wings were recorded under identical imaging conditions. Images were obtained using a Nikon SMZ 745T stereomicroscope equipped with a Labomed iVu 3100 digital camera.

Wing shape was characterized using 12 type I anatomical landmarks located at intersections between veins or between a vein and the wing margin (Fig. 1), following Prestes et al. (2023). Specimens were excluded whenever damage to either wing prevented the reliable identification of any landmark.

**Figure 1.**
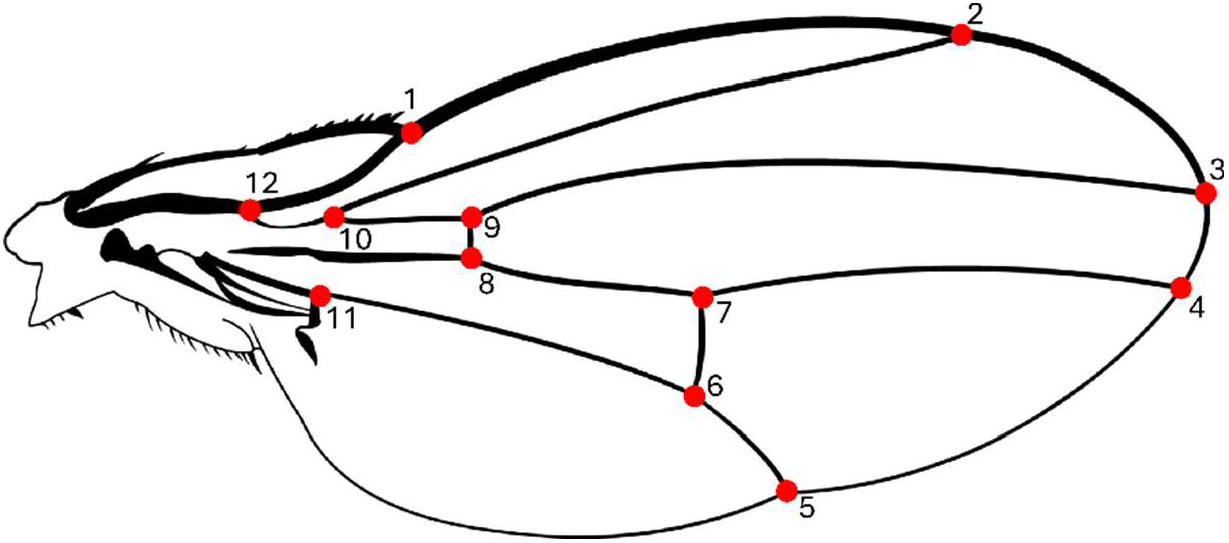
Schematic representation of the right wing of a male *Drosophila sturtevanti* in dorsal view, showing the 12 landmarks used in the geometric morphometric analyses.

Landmarks and the scale were digitized in tpsDig v.2.32 (Rohlf, 2015). The right wing was digitized first in its original orientation. The image was then horizontally mirrored, and the same landmark sequence was applied to the left wing. This procedure placed homologous landmarks of both sides in the same anatomical orientation before subsequent analyses. Each wing was digitized independently twice to quantify digitizing error. Only individuals with complete left- and right-wing data for both digitizing replicates were retained for analysis. Because left-wing images were mirrored before landmark digitization, no additional reflection of landmark coordinates was applied during subsequent shape analyses.

### 2.4. Geometric morphometric analyses

#### 2.4.1. Statistical environment and R packages

All morphometric and statistical analyses were performed in R (version 4.6.0), primarily using the packages `geomorph` (Adams *et al*., 2026; Baken *et al*., 2021) and `RRPP` (Collyer and Adams, 2018, 2024; Wickham, 2016; Wickham *et al*., 2026). Data manipulation and figure preparation were carried out using `dplyr` (Wickham *et al*., 2026), `tidyr` (Wickham *et al*., 2025), and `ggplot2` (Wickham, 2016).

#### 2.4.2. Generalized Procrustes Analysis

Landmark configurations were subjected to a Generalized Procrustes Analysis (GPA) using the `gpagen` function in the `geomorph` package. The GPA removed variation due to position, orientation, and scale, yielding Procrustes coordinates for shape analyses. Because left-wing images had already been mirrored before landmark digitization, no additional reflection was applied during the GPA.

#### 2.4.3. Digitizing error assessment

Digitizing error was assessed using the two independent landmark replicates obtained for each wing. Procrustes ANOVA was performed separately for each sampling year, partitioning variation into `individual`, `side`, `individual × side` interaction, and `residual` components. In this design, the `individual × side` interaction represents biological fluctuating asymmetry, whereas the `residual` represents digitizing error.

#### 2.4.4. Assessment of allometry

After no systematic digitizing bias was detected and random digitizing error was shown to be smaller than the individual-by-side variation, the two digitizing replicates of each wing were averaged landmark by landmark.

To assess whether wing shape covaried with size, the mean wing shape of each individual was obtained by averaging the Procrustes coordinates of the left and right wings; this configuration represents the symmetric component of wing shape. Centroid size was averaged between the left and right wings for each individual and log-transformed. Allometry was evaluated using a Procrustes linear model including sampling year, log-transformed bilateral mean centroid size, and their interaction (`shape ∼ Year * logCS`). Significance was assessed by residual randomization with 9,999 permutations. A significant effect of `logCS` was interpreted as evidence of allometry, whereas the `Year × logCS` interaction tested whether the relationship between wing shape and size differed among sampling years.

#### 2.4.5. Total Assymetry

For each individual, wing-shape asymmetry was defined as the vector difference between the Procrustes coordinates of the right and left wings (*A*_i_ = *R*_i_ − *L*_i_). Because the configurations had already been aligned by GPA, this vector represents left–right shape differences after removal of variation due to position, orientation, and scale. The magnitude of the shape asymmetry was quantified as the squared norm of this vector, obtained by summing the squared coordinate differences between homologous landmarks of the right and left wings.

For each sampling year, directional asymmetry (*DA*) was defined as the mean asymmetry vector across individuals 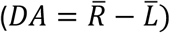. The magnitude of DA was quantified as the squared norm of this mean vector. Fluctuating asymmetry (*FA*) for each individual was defined as the deviation of that individual’s asymmetry vector from the corresponding annual *DA* vector (*FA*_i_ = *A*_i_ − *DA*). Its magnitude was quantified as the squared norm of the *FA* vector.

Because the mean *FA* vector within each year is zero by definition, total squared shape asymmetry can be decomposed exactly into directional and fluctuating components:

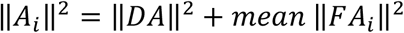

This decomposition was used to quantify the relative contribution of *DA* and *FA* to total wing-shape asymmetry in each sampling year. Differences in total squared shape asymmetry among years were evaluated using permutation tests, and pairwise *P*-values were adjusted using the Benjamini–Hochberg procedure.

##### 2.4.5.1. Fluctuating asymmetry (FA)

Differences in fluctuating asymmetry among sampling years were assessed using the individual squared FA magnitudes. A global permutation test was first used to evaluate differences among years, followed by pairwise permutation tests. Pairwise *P*-values were adjusted for multiple comparisons using the Benjamini– Hochberg procedure. All tests were based on 9,999 permutations.

##### 2.4.5.2. Directional asymmetry (DA)

Temporal variation in the magnitude of directional asymmetry was first evaluated as the norm of the annual mean asymmetry vector. Differences in *DA* magnitude among sampling years were assessed using pairwise permutation tests, and uncertainty was quantified by bootstrap resampling of individuals within each year (9,999 replicates). Bootstrap 95% confidence intervals were calculated for pairwise differences in DA magnitude.

Temporal variation in the spatial pattern of directional asymmetry was then evaluated by comparing annual DA vectors among sampling years using a Procrustes linear model with sampling year as the explanatory factor. Because each vector incorporates both magnitude and orientation in shape space, significant differences among years indicate temporal changes in the overall pattern of right– left shape differences. Pairwise differences between annual DA vectors were assessed by permutation, with *P*-values adjusted using the Benjamini–Hochberg procedure.

## 3. Results

### 3.1. Environmental conditions and sampling

Environmental data for the periods preceding and encompassing field sampling showed that 2024 was consistently warmer and had lower relative humidity than 2022 and 2026 (**Fig. 2 A,B**), conditions that may have contributed to the unusually low sampling success observed in 2024. **Figure 2C** shows the sampling dates and the number of males obtained on each occasion.

**Figure 2.**
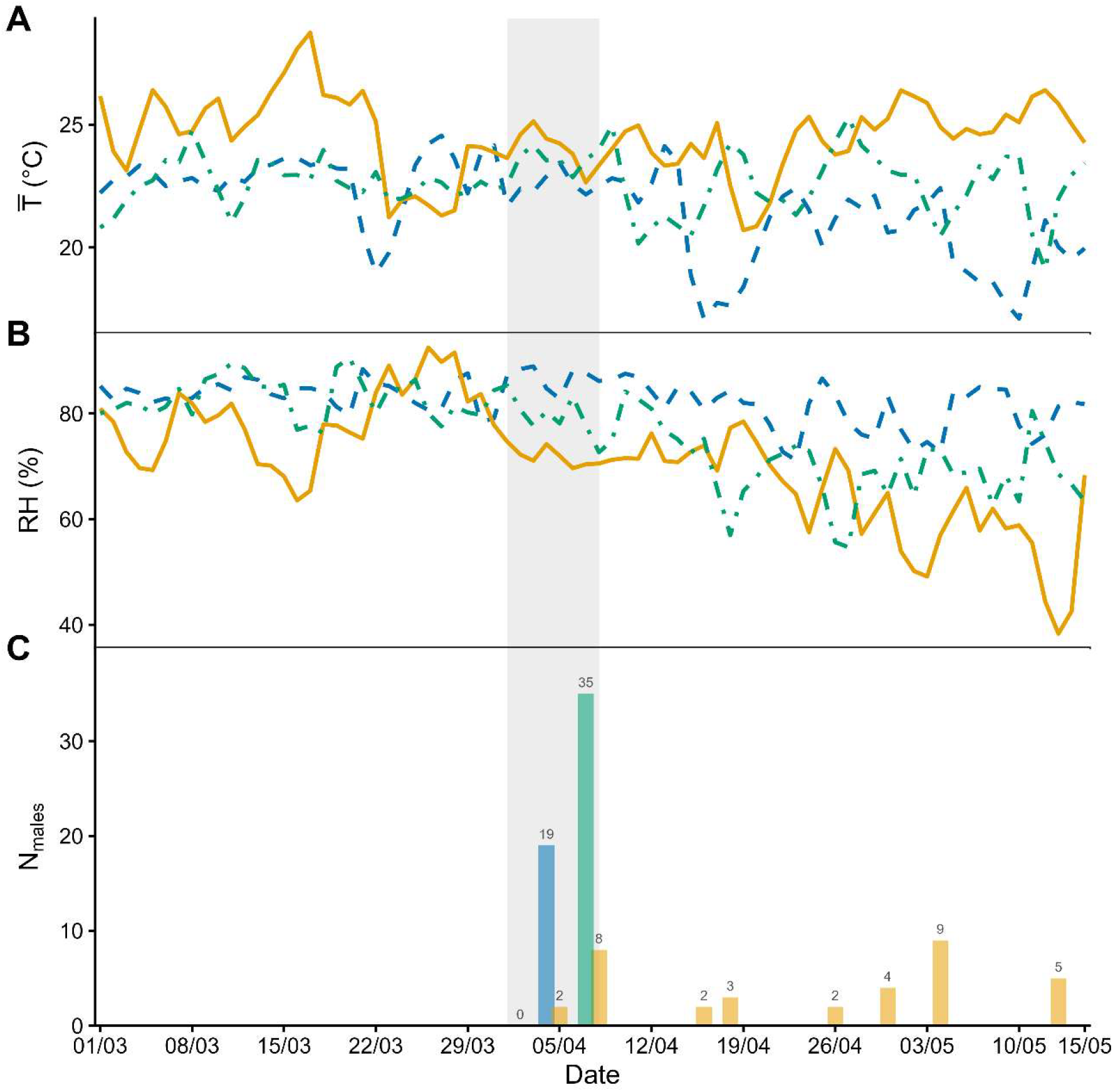
Daily mean temperature (A), daily mean relative humidity (B), and number of *Drosophila sturtevanti* males collected per sampling date (C) in the sampling locality between 1 March and 15 May in 2022, 2024, and 2026. The shaded vertical band indicates the first week of April, corresponding to the usual sampling period adopted in previous years. In panel C, a single sampling event was carried out in 2022 (*n* = 19 males) and in 2026 (*n* = 35 males), whereas repeated collections were required in 2024, yielding between 0 and 9 males per sampling date.

### 3.2. Measurement error and allometry

Wing morphology was successfully evaluated in *n* = 14, 29, and 30 males from 2022, 2024, and 2026, respectively. Fluctuating shape asymmetry exceeded digitizing error in all three sampling years. The Individual × Side effect was significant in 2022, 2024, and 2026 (all P < 0.001), with mean squares 7.7–15.0 times larger than the residual mean square (**Table S1**). Thus, digitizing error was small relative to fluctuating asymmetry and therefore unlikely to bias subsequent analyses.

No evidence of wing-shape allometry was detected. Neither log-transformed bilateral mean centroid size nor its interaction with sampling year significantly explained variation in wing shape (`logCS`: *R*^2^ = 0.0081, *F* = 0.615, *P* = 0.777; `Year × logCS`: *R*^2^ = 0.0096, *F* = 0.362, *P* = 0.9958; **Table S2**). Wing shape nevertheless differed significantly among sampling years (`Year:` *R*^2^ = 0.09453, F = 3.567, *P* < 0.001). Subsequent shape analyses were therefore performed without allometric correction.

### 3.3. Wing-shape asymmetry

Total wing-shape asymmetry was lower in 2024 than in both 2022 and 2026, whereas 2022 and 2026 did not differ detectably (**Fig. 3; Table S3**). In contrast, the magnitude of directional asymmetry remained similar across years (0.00946, 0.00887, and 0.00979 in 2022, 2024, and 2026, respectively), although directional asymmetry was not statistically detectable within the 2022 sample (**Table S4**).

**Figure 3.**
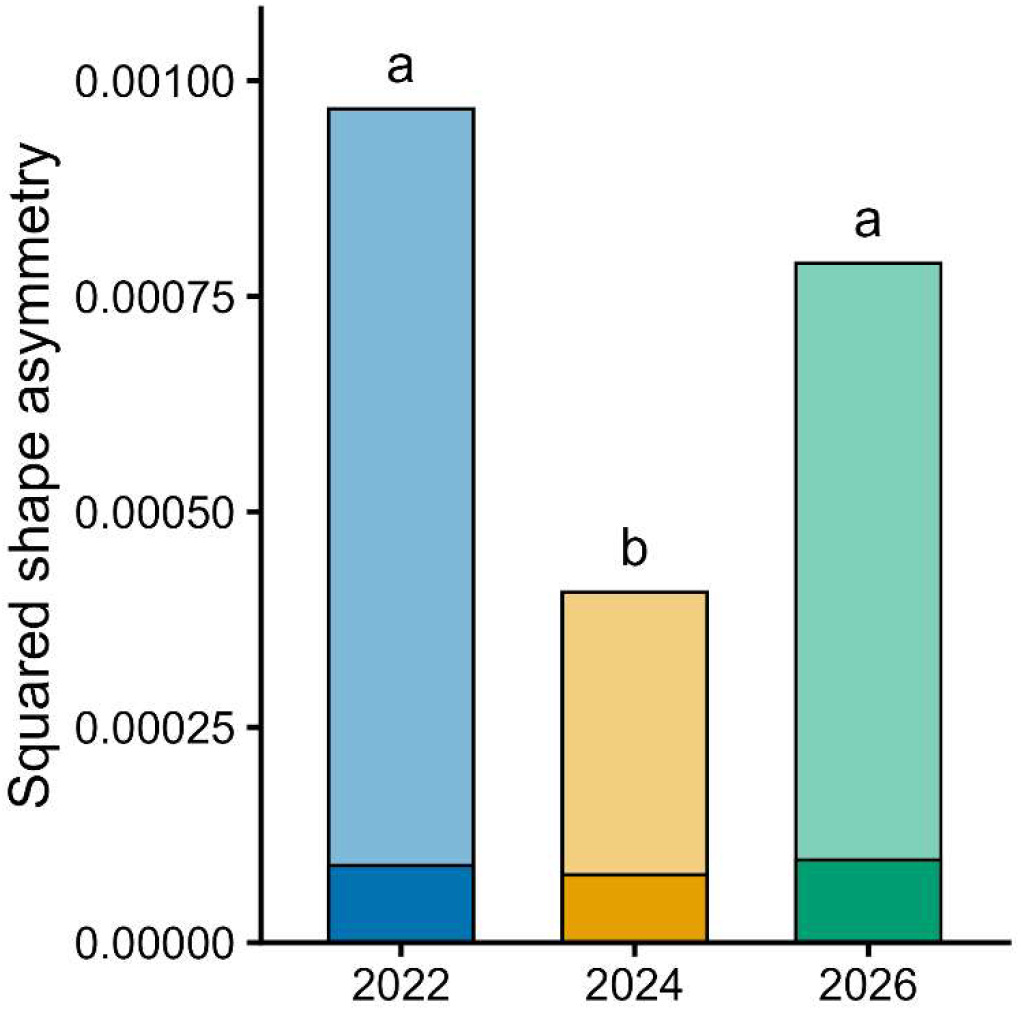
Total wing-shape asymmetry in males sampled in 2022, 2024, and 2026. The darker lower portion of each bar represents directional asymmetry (DA), whereas the lighter upper portion represents fluctuating asymmetry (FA). Lowercase letters indicate significant differences in total asymmetry among years after Benjamini–Hochberg correction (see **Table S3**).

Pairwise permutation tests detected no differences in DA magnitude among years after BH correction, and all bootstrap 95% confidence intervals for pairwise differences included zero (**Table S5**). Thus, temporal differences in total asymmetry were driven primarily by variation in fluctuating asymmetry, which was lower in 2024 than in both 2022 and 2026 (**Fig. 4; Table S6**).

**Figure 4.**
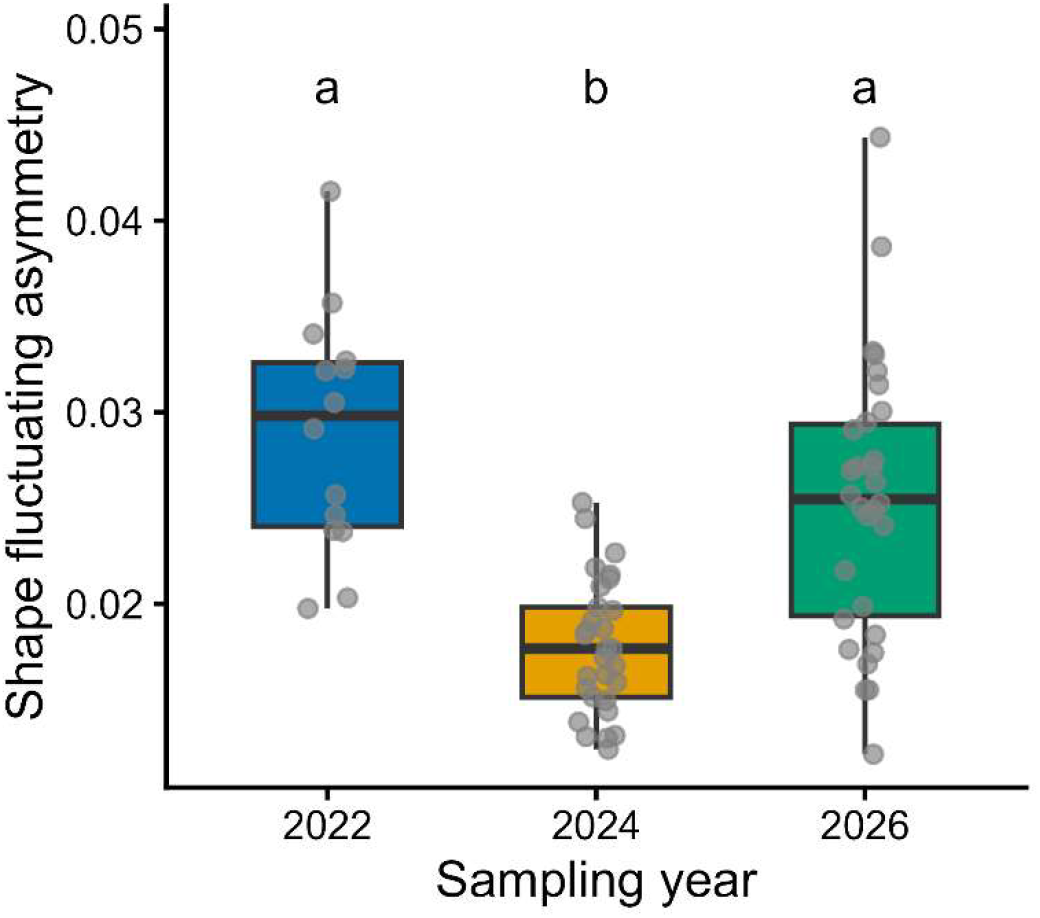
Fluctuating asymmetry of wing shape in males sampled in 2022, 2024, and 2026. Boxplots show the distribution of individual fluctuating-asymmetry magnitudes for each sampling year; points represent individual males. Lowercase letters indicate significant differences among years after Benjamini–Hochberg correction (see **Table S6**).

Although DA magnitude did not differ among years, the spatial pattern of directional asymmetry varied temporally. The 2022 DA vector did not differ detectably from either 2024 or 2026, whereas 2024 and 2026 differed significantly (**Fig. 5; Table 1**).

**Table 1.** Pairwise comparisons of directional asymmetry vectors among sampling years.

| Comparison | N1 | N2 | DA vector distance | P_BH |
| --- | --- | --- | --- | --- |
| 2022 vs 2024 | 14 | 29 | 0.00626 | 0.732 |
| 2022 vs 2026 | 14 | 30 | 0.0128 | 0.0554 |
| 2024 vs 2026 | 29 | 30 | 0.0135 | 0.0006 |

**Note:** After Benjamini–Hochberg correction, only the 2024 vs 2026 comparison was significant.

**Figure 5.**
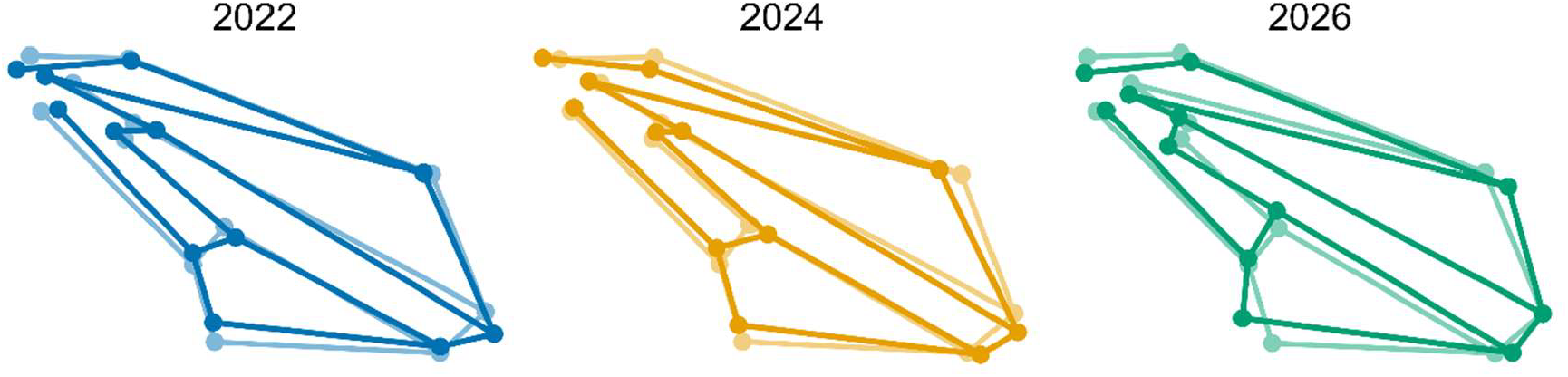
Directional asymmetry in wing shape of males sampled in 2022, 2024, and 2026. For each sampling year, the mean configurations of the left and right wings are shown superimposed. The lighter configuration represents the mean left wing, whereas the darker configuration represents the mean right wing. Differences between sides were magnified 10× to facilitate visualization of the spatial pattern of directional asymmetry.

## 4. Discussion

The unusually low sampling success of *D. sturtevanti* in April 2024, compared with 2022 and 2026, prompted us to ask whether that year was characterized by atypical environmental conditions. NASA POWER data showed that, throughout March and April, 2024 was consistently warmer and had lower relative humidity than the other two sampling years, a pattern that persisted until mid-May, when sampling was terminated. Although this association is strictly correlative, high temperatures and reduced humidity may affect food availability as well as survival and reproductive conditions in the field, making it plausible that the low sampling success observed in 2024 was related, at least in part, to the environmental conditions prevailing that year.

El Niño is a recurrent climatic phenomenon characterized by anomalous warming of the tropical Pacific Ocean and associated with widespread effects on weather patterns across the globe(Cai *et al*., 2021). One such event occurred between 2023 and 2024 and provides a broader climatic context for the pattern observed here. This event was exceptionally strong, ranking as the fourth strongest event since 1979 (Zhang *et al*., 2025)and reaching its peak between November 2023 and January 2024 (WMO, 2025). Although it subsequently weakened, its influence on global climate persisted into the following months, and El Niño conditions were still present during the first months of 2024 (WMO, 2024). In Brazil, official monitoring in April 2024 likewise indicated that the event was weakening but had not yet fully transitioned to neutral conditions, with neutral conditions expected during the April–June period (ANA, 2024). Thus, our April 2024 sampling took place during the final phase of an unusually strong El Niño event.

Our second question concerned whether the environmental differences observed among years were associated with changes in wing asymmetry. Surprisingly, total asymmetry was significantly lower in 2024. When asymmetry was decomposed into directional and fluctuating components, however, contrasting temporal patterns emerged. The magnitude of directional asymmetry remained remarkably similar across years, with no significant differences among 2022, 2024, and 2026. This result is consistent with the traditional view of DA as a relatively conserved feature of populations or species, reflecting consis(Klingenberg *et al*., 1998)ed development (Klingenberg *et al*., 1998). Nevertheless, DA is not necessarily invariant. Costa et al. (2015), for example, found geographic differences in DA magnitude among populations of *Drosophila antonietae* Tidon-Sklorz and Sene, 2001, whereas FA remained comparatively constant across the species’ distribution.

Our results reveal another dimension of this variation. Although DA magnitude remained stable through time, the spatial pattern of the DA vector differed significantly between 2024 and 2026, while neither differed detectably from 2022. In geometric morphometrics, vector length describes the magnitude of shape asymmetry, whereas differences in vector direction reflect differences in the spatial pattern of asymmetry (Mitteroecker and Schaefer, 2022). Thus, temporal stability in the magnitude of DA did not imply stability in how directional asymmetry was expressed across the wing. Together with the geographic variation reported by Costa et al. (2015), this suggests that DA may be more labile within species than implied by its traditional characterization as a fixed population or species-level attribute.

The reduction in total asymmetry observed in 2024 was therefore primarily attributable to a reduction in FA. This result was unexpected, since environmental stress has generally been associated with increased fluctuating asymmetry (Beasley *et al*., 2013). Because the two sides of a bilaterally symmetrical structure share the same genotype and develop under broadly similar environmental conditions, random differences between sides have traditionally been interpreted as manifestations of developmental instability (Klingenberg and McIntyre, 1998). Thus, given the unusually warm and dry conditions observed in 2024, an increase rather than a reduction in FA might have been expected.

Nevertheless, the relationship between environmental stress and FA is not necessarily monotonic. Experimental studies show that severe stress can substantially increase mortality without producing the expected increase in FA. Polak et al. (2002), for example, exposed developing *Drosophila melanogaster* Meigen, 1830 to arsenic and showed that wing FA covaried positively with adult emergence, with lower emergence associated with lower FA. They interpreted this pattern as evidence of developmental selection against less developmentally stable individuals. Similarly, Floate & Fox (2000) found sharply reduced pupation success under increasing ivermectin exposure in *Musca domestica*, but no corresponding change in wing FA, and suggested that differential mortality may leave a more robust subset among the survivors.

A similar process could provide one explanation for our field observations. The unusually warm and dry conditions of 2024 coincided with both low sampling success and reduced FA among the adults collected. If individuals expressing greater developmental instability were less likely to survive to adulthood or, after emergence, were less able to persist in the field and reach banana-baited traps, the sampled adults could represent a selectively filtered subset of the population, consistent with evidence that more asymmetrical individuals may show reduced survival and competitive performance (Møller, 1997). This interpretation remains hypothetical because mortality and developmental success were not measured directly in the field, but it provides a plausible explanation for why FA decreased rather than increased under apparently more challenging conditions.

More broadly, the relationship between environmental adversity and FA is not straightforward. Empirical studies have reported positive, absent, and even inverse associat(Graham, 2021; Lens *et al*., 2002) and FA (Graham, 2021; Lens *et al*., 2002). Symanski & Redak (2021), for instance, found that increased stress was associated with higher mortality but no detectable change in FA in Lepidoptera. Particularly relevant to our study, Nascimento et al. (2021) tested whether *D. sturtevanti* and other drosophilid species from the Brazilian Cerrado would exhibit greater FA during the dry season, when environmental conditions are generally considered less favorable. Contrary to this expectation, they found no significant difference in FA between the dry and rainy seasons. The authors suggested that increased mortality during the drier season could contribute to maintaining similar levels of asymmetry among the surviving adults.

Our study differs from this seasonal context because the unusually warm and dry conditions of 2024 occurred during a period in which *D. sturtevanti* is normally abundant. This contrast raises a broader question: how do FA and population abundance respond when environmental conditions depart substantially from those normally experienced by a population at a given time of year? If such conditions impose strong demographic filtering, as suggested by experimental studies in which arsenic or ivermectin exposure caused high developmental mortality without the expected increase in FA (Floate and Fox, 2000; Polak *et al*., 2002), the increasing frequency of extreme climatic events makes it increasingly important to understand how organisms respond to abrupt environmental departures from typical seasonal conditions. In our system, this leads to a clear prediction: if sampling is repeated at the same site and during the same seasonal window under the influence of a comparable extreme climatic event, both reduced capture success and reduced FA should again be observed among the adults collected.

The ongoing 2026–2027 El Niño, which is currently predicted to persist into the first months of 2027 (CPC, 2026a; 2026b), provides an immediate opportunity to test this prediction in our *D. sturtevanti* population. More broadly, it may offer a valuable natural experiment for researchers with previous field samples from fixed sites, allowing comparisons of abundance and FA before and during an unusually strong climatic event.

## Acknowledgments

We thank Marina Magalhães Moreira, Luísa de Paula Bouzada Dias, and Letícia Carlesso de Paula Sena for substantial assistance with field sampling in 2022 and 2024. Marina also contributed to specimen sorting in both years, and Luísa photographed the 2022 wings. We also thank Letícia de Castro Araújo for assistance with field sampling in 2026. We acknowledge the Coordenação de Aperfeiçoamento de Pessoal de Nível Superior (CAPES) for institutional support, the Graduate Program in Ecology and the Department of General Biology at Universidade Federal de Viçosa for infrastructure and logisti cal support, and the Fundação de Amparo à Pesquisa do Estado de Minas Gerais (FAPEMIG) for the scholarship awarded to Caio de Castro Mello (14457) and Pedro Gabriel Carneiro Moreira (16026).

## References

Adams, D.C., Collyer, M.L., Kaliontzopoulou, A. and Baken, E.K. (2026), “Geomorph: Software for geometric morphometric analyses. R package version”, CRAN.

Agência Nacional de Águas e Saneamento Básico (ANA). (2024), “Última edição do boletim sobre El Niño indica que fenômeno tende a terminar entre abril e junho”, Agência Nacional de Águas e Saneamento Básico, available at: https://www.gov.br/ana/pt-br/assuntos/noticias-e-eventos/noticias/ultima-edicao-do-boletim-sobre-el-nino-indica-que-fenomeno-tende-a-terminar-entre-abril-e-junho x(accessed 22 August 2026).

Baken, E.K., Collyer, M.L., Kaliontzopoulou, A. and Adams, D.C. (2021), “geomorph v4.0 and gmShiny: Enhanced analytics and a new graphical interface for a comprehensive morphometric experience”, Methods in Ecology and Evolution, Vol. 12 No. 12, pp. 2355–2363, doi: 10.1111/2041-210X.13723.

Beasley, D.A.E., Bonisoli-Alquati, A. and Mousseau, T.A. (2013), “The use of fluctuating asymmetry as a measure of environmentally induced developmental instability: A meta-analysis”, Ecological Indicators, Vol. 30, pp. 218–226, doi: 10.1016/j.ecolind.2013.02.024.

Cai, W., Santoso, A., Collins, M., Dewitte, B., Karamperidou, C., Kug, J.S., Lengaigne, M., et al. (2021), “Changing El Niño–Southern Oscillation in a warming climate”, Nature Reviews Earth and Environment, Springer Nature, 1 September, doi: 10.1038/s43017-021-00199-z.

Climate Prediction Center (CPC). (2026a), “Official NOAA CPC ENSO Probabilities”, National Centers for Environmental Prediction, available at: https://cpc.ncep.noaa.gov/products/analysis_monitoring/enso/roni/probabil ities/(accessed 22 August 2026).

Climate Prediction Center (CPC). (2026b), “El Niño/Southern Oscillation (ENSO) Diagnostic Discussion”, National Oceanic and Atmospheric Administration, available at: https://www.cpc.ncep.noaa.gov/products/analysis_monitoring/enso_advisory/ensodisc.html (accessed 22 August 2026).

Collyer, M.L. and Adams, D.C. (2018), “RRPP: An R package for fitting linear models to high-dimensional data using residual randomization”, Methods in Ecology and Evolution, Vol. 9 No. 7, pp. 1772–1779, doi: 10.1111/2041-210X.13029.

Collyer, M.L. and Adams, D.C. (2024), “Linear Model Evaluation with Randomized Residuals in a Permutation”, CRAN.

Costa, M., Mateus, R.P. and Moura, M.O. (2015), “Constant fluctuating asymmetry but not directional asymmetry along the geographic distribution of Drosophila antonietae (Diptera, Drosophilidae)”, Revista Brasileira de Entomologia, Sociedade Brasileira De Entomologia, Vol. 59 No. 4, pp. 337– 342, doi: 10.1016/j.rbe.2015.09.004.

Dias, L. de P.B., Moreira, M.M., Sena, L.C. de P. and Yotoko, K.S.C. (2026), “Rapid changes in wing size and shape during the transition to laboratory conditions in a Neotropical drosophilid”, BioRxiv, doi: 10.64898/2026.09.03.749121.

Duda, O. (1927), “Die südamerikanischen Drosophiliden (Dipteren) unter Berücksichtigung auch der anderen neotropischen sowie der nearktischen Arten”, Archiv Für Naturgeschichte, Vol. 91 No. 11/12, pp. 1–228.

Floate, K.D. and Fox, A.S. (2000), “Flies under stress: A test of fluctuating asymmetry as a biomonitor of environmental quality”, Ecological Applications, Ecological Society of America, Vol. 10 No. 5, pp. 1541–1550, doi: 10.1890/1051-0761(2000)010[1541:FUSATO]2.0.CO;2.

Galego, L.G.C. and Carareto, C.M.A. (2005), “Intraspecific and interspecific pre-adult competition on the Neotropical region colonizer Zaprionus indianus (Diptera: Drosophilidae) under laboratory conditions”, Bragantia, Vol. 64, pp. 249–255.

Graham, J.H. (2021), “Nature, Nurture, and Noise: Developmental Instability, Fluctuating Asymmetry, and the Causes of Phenotypic Variation”, Symmetry, MDPI AG, Vol. 13 No. 7, p. 1204, doi: 10.3390/sym13071204.

Klingenberg, C.P. (2022), “Shape asymmetry — what’s new?”, Emerging Topics in Life Sciences, Portland Press Ltd, Vol. 6 No. 3, pp. 285–294, doi: 10.1042/ETLS20210273.

Klingenberg, C.P. and McIntyre, G.S. (1998), “Geometric morphometrics of developmental instability: Analyzing patterns of fluctuating asymmetry with procrustes methods”, Evolution, Society for the Study of Evolution, Vol. 52 No. 5, pp. 1363–1375, doi: 10.1111/j.1558-5646.1998.tb02018.x.

Klingenberg, C.P., McIntyre, G.S. and Zaklan, S.D. (1998), “Left-right asymmetry of fly wings and the evolution of body axes”, Proceedings of the Royal Society of London. Series B: Biological Sciences, pp. 1255–1259.

Lens, L., van Dongen, S., Kark, S. and Matthsen, E. (2002), “Fluctuating asymmetry as an indicator of fitness: can we bridge the gap between studieş”, Biological Reviews, Cambridge Philosophical Society, Vol. 77 No. 1, pp. 27–38, doi: 10.1017/S1464793101005796.

Magalhães, L. (1962), “Notes on the taxonomy, morphology, and distribution of the Saltans group of Drosophila, with description of four new species”, The University of Texas Publication, Vol. 6205, pp. 135–154.

Medeiros, H.F. and Klaczko, L.B. (1999), “A weakly biased Drosophila trap”, Drosophila Information Service, Vol. 82, pp. 100–102.

Mitteroecker, P. and Schaefer, K. (2022), “Thirty years of geometric morphometrics: Achievements, challenges, and the ongoing quest for biological meaningfulness”, American Journal of Biological Anthropology, John Wiley and Sons Inc, Vol. 178 No. S74, pp. 181–210, doi: 10.1002/ajpa.24531.

Møller, A.P. (1997), “Developmental Stability and Fitness: A Review”, The American Naturalist, Vol. 149 No. 5, pp. 916–932, doi: 10.1086/286030.

Nascimento, L.C., Machado, F. das C.R. and Tidon, R. (2021), “Drosophilid species (Insecta, Diptera) in the Brazilian savanna are more symmetric in the dry season”, Heringeriana, Jardim Botânico de Brasília, Vol. 15, pp. 17–26, doi: 10.17648/heringeriana.v15i1.917941.

Ooms, J. (2014), “The jsonlite Package: A Practical and Consistent Mapping Between JSON Data and R Objects”, ArXiv, p. 1403.2805.

Palmer, A.R. and Strobeck, C. (1986), “Fluctuating asymmetry: measurement, analysis, patterns”, Annual Review of Ecology and Systematics, Vol. 17, pp. 391–421.

Pedersen, T.L. (2019), “patchwork: The Composer of Plots”, CRAN: Contributed Packages, 1 December, doi: 10.32614/CRAN.package.patchwork.

Polak, M., Opoka, R. and Cartwright, I.L. (2002), “Response of ?uctuating asymmetry to arsenic toxicity: support for the developmental selection hypothesis”, Environmental Pollution, Vol. 118 No. 1, pp. 19–28, doi: 10.1016/S0269-7491(01)00281-0.

Prestes, J.O., Costa, M., Machado, L.P.B. and Mateus, R.P. (2023), “Asymmetry of the wing in Drosophila mojavensis sonorensis Castrezana in Pfeiler et al., 2009 (Diptera, Drosophilidae): main versus seasonal host”, Brazilian Journal of Biology, Instituto Internacional de Ecologia, Vol. 83, p. e246677, doi: 10.1590/1519-6984.246677.

Rohlf, F.J. (2015), “The tps series of software”, Hystrix, Associazione Teriologica Italiana onlus, Vol. 26 No. 1, pp. 1–4, doi: 10.4404/hystrix-26.1-11264.

Symanski, C. and Redak, R.A. (2021), “Does fluctuating asymmetry of wing traits capture relative environmental stress in a lepidopteran?”, Ecology and Evolution, John Wiley and Sons Ltd, Vol. 11 No. 3, pp. 1199–1213, doi: 10.1002/ece3.7097.

Thompson, D.W. (1994), On Growth and Form, edited by Bonner, J.T., Cambridge University Press, Cambridge, UK.

Tidon-Sklorz, R. and Sene, F.M. (2001), “Two new species of the Drosophila serido sibling set (Diptera, Drosophilidae)”, Iheringia, Série Zoologia, Vol. 90, pp. 141–146.

Wickham, H. (2016), Ggplot2: Elegant Graphics for Data Analysis, 2nd ed., Springer International Publishing, Cham, doi: 10.1007/978-3-319-24277-4.

Wickham, H., François, R., Henry, L., Müller, K. and Vaughan, D. (2026), “dplyr: A Grammar of Data Manipulation”, CRAN: Contributed Packages, CRAN, doi: 10.32614/CRAN.package.dplyr.

Wickham, H., Vaughan, D. and Girlich, M. (2025), “tidyr: Tidy Messy Data”, CRAN: Contributed Packages, doi: 10.32614/CRAN.package.tidyr.

World Meteorological Organization (WMO). (2024), “El Niño weakens but impacts continue”, available at: https://wmo.int/news/media-centre/el-nino-weakens-impacts-continue x(accessed 22 August 2026).

World Meteorological Organization (WMO). (2025), State of the Global Climate 2024, Geneva, Switzerland.

Zelditch, M.L., Swiderski, D.L., Sheets, H.D. and Fink, W.L. (2012), Geometric Morphometrics for Biologists : A Primer, 2nd ed., Elsevier Academic Press.

Zhang, L., Chen, Y., Karnauskas, K.B., Wang, C., Collins, M. and Luo, X. (2025), “The 2023/24 El Niño event exhibited unusually weak extratropical teleconnections”, Communications Earth and Environment, Nature Publishing Group, Vol. 6 No. 1, doi: 10.1038/s43247-025-02584-8.

