## Supplemental Material for "El Niño, capture success, and wing fluctuating asymmetry in *Drosophila sturtevanti* Duda, 1927"

### Supplementar Material

**Table S1. Procrustes ANOVA results used to assess digitizing error in wing shape for males sampled in 2022, 2024, and 2026.** Mean squares are shown for the Individual × Side interaction and the residual component.

| Year | MS Individual × Side | MS residual | $F$ | $P$ |
|---|---|---|---|---|
| 2022 | 0.00094587 | 0.00007 | 13.513 | <0.001 |
| 2024 | 0.00033978 | 0.00004419 | 7.689 | <0.001 |
| 2026 | 0.00071691 | 0.00004781 | 14.996 | <0.001 |

**Note:** The Individual × Side interaction represents fluctuating asymmetry, whereas the residual component contains digitizing error together with any other unexplained variation. $P$-values refer to the Individual × Side effect, and the reported $F$ statistic is the ratio between the mean square of this interaction and the residual mean square.

**Table S2:** Procrustes linear model used to assess wing-shape allometry across sampling years.

| Effect | Df | SS | MS | $R^2$ | F | Z | $P$ |
|---|---|---|---|---|---|---|---|
| Year | 2 | 0.002984 | 0.001492 | 0.09453 | 3.567 | 4.2775 | <0.001 |
| logCS | 1 | 0.000257 | 0.000257 | 0.00814 | 0.6146 | -0.7731 | 0.7773 |
| Year × logCS | 2 | 0.000303 | 0.000151 | 0.00958 | 0.3616 | -2.6276 | 0.9958 |
| Residuals | 67 | 0.028024 | 0.000418 | 0.88775 | | | |

**Note:** Neither logCS nor the Year × logCS interaction significantly explained variation in wing shape ($P > 0.05$). Therefore, subsequent shape analyses were performed without allometric correction.

**Table S3.** Pairwise comparisons of total squared wing-shape asymmetry among sampling years. Mean values refer to total squared wing-shape asymmetry.

| Comparison | Mean 1 | Mean 2 | Difference | Ratio | $P$_BH |
|---|---|---|---|---|---|
| 2022 vs 2024 | 0.000967 | 0.000407 | 0.000561 | 2.38 | 0.00015 |
| 2022 vs 2026 | 0.000967 | 0.000788 | 0.000179 | 1.23 | 0.222 |
| 2024 vs 2026 | 0.000407 | 0.000788 | -0.000382 | 0.516 | 0.00015 |

**Note:** Total squared wing-shape asymmetry was lower in 2024 than in both 2022 and 2026 after Benjamini–Hochberg correction, whereas 2022 and 2026 did not differ significantly.

**Table S4**. Directional asymmetry within each sampling year.

| Year | $N$ | DA magnitude | $P$ | $P$_BH |
|---|---|---|---|---|
| 2022 | 14 | 0.00946 | 0.213 | 0.213 |
| 2024 | 29 | 0.00887 | 0.0001 | 0.0003 |
| 2026 | 30 | 0.00979 | 0.0009 | 0.00135 |

**Note:** Directional asymmetry was statistically detectable in 2024 and 2026, but not in 2022.

**Table S5.** Pairwise comparisons of directional asymmetry (DA) magnitude among sampling Years.

| Comparison | Magnitude 1 | Magnitude 2 | Difference | ↓95% CI | ↑95% CI | $P$_BH |
|---|---|---|---|---|---|---|
| 2022 vs 2024 | 0.00946 | 0.00887 | 0.00059 | -0.00124 | 0.00761 | 0.907 |
| 2022 vs 2026 | 0.00946 | 0.00979 | -0.000327 | -0.0034 | 0.00672 | 0.907 |
| 2024 vs 2026 | 0.00887 | 0.00979 | -0.000917 | -0.00543 | 0.00235 | 0.907 |

**Note:** Differences correspond to DA magnitude in the first year minus DA magnitude in the second year. Confidence intervals were obtained by bootstrap resampling with 9,999 replicates. *$P$-values were obtained by pairwise permutation tests and adjusted for multiple comparisons using the Benjamini–Hochberg procedure.*

**Table S6.** Pairwise comparisons of fluctuating wing-shape asymmetry (FA) among sampling years.

| Comparison | N1 | N2 | Mean squared FA 1 | Mean squared FA 2 | Difference | Ratio | $P$_BH |
|---|---|---|---|---|---|---|---|
| 2022 vs 2024 | 14 | 29 | 0.000878 | 0.000328 | 0.00055 | 2.68 | 0.0002 |
| 2022 vs 2026 | 14 | 30 | 0.000878 | 0.000692 | 0.000185 | 1.27 | 0.142 |
| 2024 vs 2026 | 29 | 30 | 0.000328 | 0.000692 | -0.000364 | 0.47 | 0.0002 |

**Note:** Differences correspond to mean squared FA in the first year minus mean squared FA in the second year. Ratios correspond to mean squared FA in the first year divided by that in the second year. $P$-values were obtained by permutation tests and adjusted for multiple comparisons using the Benjamini–Hochberg procedure.
